# *Pseudomonas zeiradicis* from corn and *Pseudomonas soyae* from soybean, two new *Pseudomonas* species and endophytes from agricultural crops

**DOI:** 10.1101/2021.11.02.466922

**Authors:** Jacqueline Lemaire, Sarah Seaton, Patrik Inderbitzin, Martha E. Trujillo

## Abstract

Two novel *Pseudomonas* species associated with healthy plants and other habitats are described from the United States. They are *Pseudomonas zeiradicis* strain PI116 from corn in Missouri, compost from Massachusetts, urban soil from Iowa and water of Lake Erie; and *Pseudomonas soyae* strain JL117 from soybean in Indiana and Wisconsin, and soil in Wyoming. No plant pathogenic strains are known for any of the novel species based on genome comparisons to assemblies in GenBank.

## INTRODUCTION

*Pseudomonas* is a large and diverse genus in the *Gammaproteobacteria* comprising more than 180 named species (Hesse et al. 2018). *Pseudomonas* species are rod-shaped, Gram-type negative chemoheterotrophs, which are motile by means of polar flagella, and can utilize an array of small organic molecules as sources of carbon and energy. This nutritional diversity is reflected in the high abundance of *Pseudomonas* species in nature, with as many as 10^6^ fluorescent pseudomonads residing in a single gram of soil (Tarnawski et al. 2003; Vančura 1980). *Pseudomonas* species are strictly aerobic, but some can utilize NO_3_ as an electron acceptor in place of O_2_. In addition, nitrate can be used as a nitrogen source for all known species (Stanier et al. 1966).

Pseudomonads are ubiquitous in the rhizosphere, often living in a commensal relationship with plants and utilizing plant-exuded nutrients. *Pseudomonas* species also impact plant health by suppressing phytopathogens (Couillerot et al. 2009; Gerbore et al. 2014), enhancing local access to nutrients, and inducing systemic resistance in the plant host (Bakker et al. 2007; De Vleesschauwer et al. 2008). A subset of rhizosphere pseudomonads are further adapted to life as endophytes, residing within cells, the intercellular spaces or the vascular system of host plants (Mitter et al. 2013). We describe two such endophytic strains, representing two novel species of the genus *Pseudomonas. Pseudomonas zeiradicis* sp. nov. strain PI116, which was isolated from healthy, field-grown corn in Missouri, United States, and *Pseudomonas soyae* sp. nov. strain JL117 from healthy, field-grown soybean in Indiana, United States. We provide phenotypic and phylogenomic details describing these two species, and report geographic distribution and additional substrates from conspecific, cultured strains by comparison to whole genomes at GenBank.

## METHODS

### Isolation

Strain PI116 was isolated from healthy field-grown *Zea mays* roots in Missouri, United States, and strain JL117 from seedlings of *Glycine max* in Indiana, United States. Plant tissue was washed with a mild detergent to remove particulates, surface-sterilized with bleach (1% v/v sodium hypochlorite) and ethanol (70% v/v), and homogenized. Serial dilutions of tissue homogenate were plated on a panel of media types for endophyte cultivation. All strains were streaked to purity and stored in glycerol (20% v/v) at -80°C until subjected to further testing.

### Motility

Strains were tested for flagellar-dependent swimming and swarming motility on R2A plates solidified with 0.3% and 0.6% agar, respectively. Three independent colonies were inoculated onto R2A broth and grown for 36 hr at 24°C. Broth cultures were normalized to an OD600 of 0.1, and 1.5 µl of culture was spotted directly onto the surface of the motility agar. The diameter of colony expansion was measured for 5 days.

### Carbon source utilization

Substrate utilization was assessed using Biolog GenIII Microplates (Catalogue No. 1030) (Biolog Inc., Hayward, CA). Each bacterium was inoculated in duplicate plates using Protocol A described by the manufacturer, with the exception that plates were incubated at 30°C for 36 hr. Respiration leading to reduction of the tetrazolium indicator was measured by absorbance at 590 nm.

### Biochemical analyses

Catalase activity was evaluated by immediate effervescence after the application of 3 % (v/v) hydrogen peroxide solution via the tube method, a positive reaction was indicated by the production of bubbles. *Staphylococcus aureus* NCIMB 12702 and *Streptococcus pyogenes* ATCC 19615 were used as positive and negative controls, respectively. Oxidase activity was evaluated via the oxidation of Kovács oxidase reagent, 1% (w/v) tetra-methyl-p-phenylenediamine dihydrochloride in water, via the filter-paper spot method. A positive reaction was indicated when the microorganism’s color changed to dark purple. *Pseudomonas aeruginosa* NCIMB 12469 and *Escherichia coli* ATCC 25922 were used as positive and negative controls, respectively.

### Phylogenetic and genomic analyses

DNA was extracted from pure cultures using the Omega Mag-Bind Universal Pathogen Kit according to manufacturer’s protocol with a final elution volume of 60µl (Omega Biotek Inc., Norcross, GA). DNA samples were quantified using Qubit fluorometer (ThermoFisher Scientific, Waltham, MA) and normalized to 100 ng. DNA was prepped using Nextera DNA Flex Library Prep kit according to manufacturer’s instructions (Illumina Inc., San Diego, CA). DNA libraries were quantified via qPCR using KAPA Library Quantification kit (Roche Sequencing and Life Science, Wilmington, MA) and combined in equimolar concentrations into one 24-sample pool. Libraries were sequenced on a MiSeq using pair-end reads (2×200bp). Reads were trimmed of adapters and low-quality bases using Cutadapt (version 1.9.1) and assembled into contigs using MEGAHIT (version 1.1.2) (Li et al. 2015). Reads were mapped to contigs using Bowtie2 (version 2.3.4) (Langmead and Salzberg 2012), and contigs were assembled into scaffolds using BESST (2.2.8) (Sahlin et al. 2014).

16S rRNA gene sequences were extracted from genome assemblies using barrnap (Seemann 2019) and submitted to GenBank (OL306320, OL306321).

Phylogenomic analyses were performed using GToTree (Lee 2019) with default settings. Taxon sampling for each species is described in the respective phylogenetic tree figure legend.

Average nucleotide identity analyses were performed using the pyani ANIm algorithm (Richter and Rosselló-Móra 2009) implemented in the MUMmer package (Kurtz et al. 2004) retrieved from https://github.com/widdowquinn/pyani.

Geographic distribution and host range of novel species were inferred by ANI to assemblies from unidentified strains from GenBank (Ciufo et al. 2018) and the Indigo internal collection. An ANI threshold of ≥95% indicated conspecificity (Chun et al. 2018; Richter and Rosselló-Móra 2009). Distribution maps were generated in R version 4.0.4 (R Core Team 2021) with tigris 1.5 (Walker 2021), sf 1.0-2 (Pebesma 2018), ggplot2 3.3.2 (Wickham 2016), cowplot 1.1.1 (Wilke 2020) and other packages.

## RESULTS

### Genomic analyses

*Pseudomonas zeiradicis* sp. nov. strain PI116

Strain PI116 shared 99.9% 16S rRNA gene sequence identity with *Pseudomonas mohnii* CCUG 53115^T^ and less with the remaining *Pseudomonas* species. A phylogenomic tree using GToTree (Lee 2019) confirmed the affiliation of strain PI116 with the genus *Pseudomonas*. PI116 was most closely related to *P. umsongensis* DSM 16611^T^ with 100% bootstrap support (Figure 1). The top average nucleotide identity (ANI) value of PI116 was 90.8% with *P. umsongensis*. This value was well below the threshold for species demarcation (Richter and Rosselló-Móra 2009; Chun et al. 2018) providing further genomic support that strain PI116 represents a new genomic species of *Pseudomonas*.

**Figure 1.**
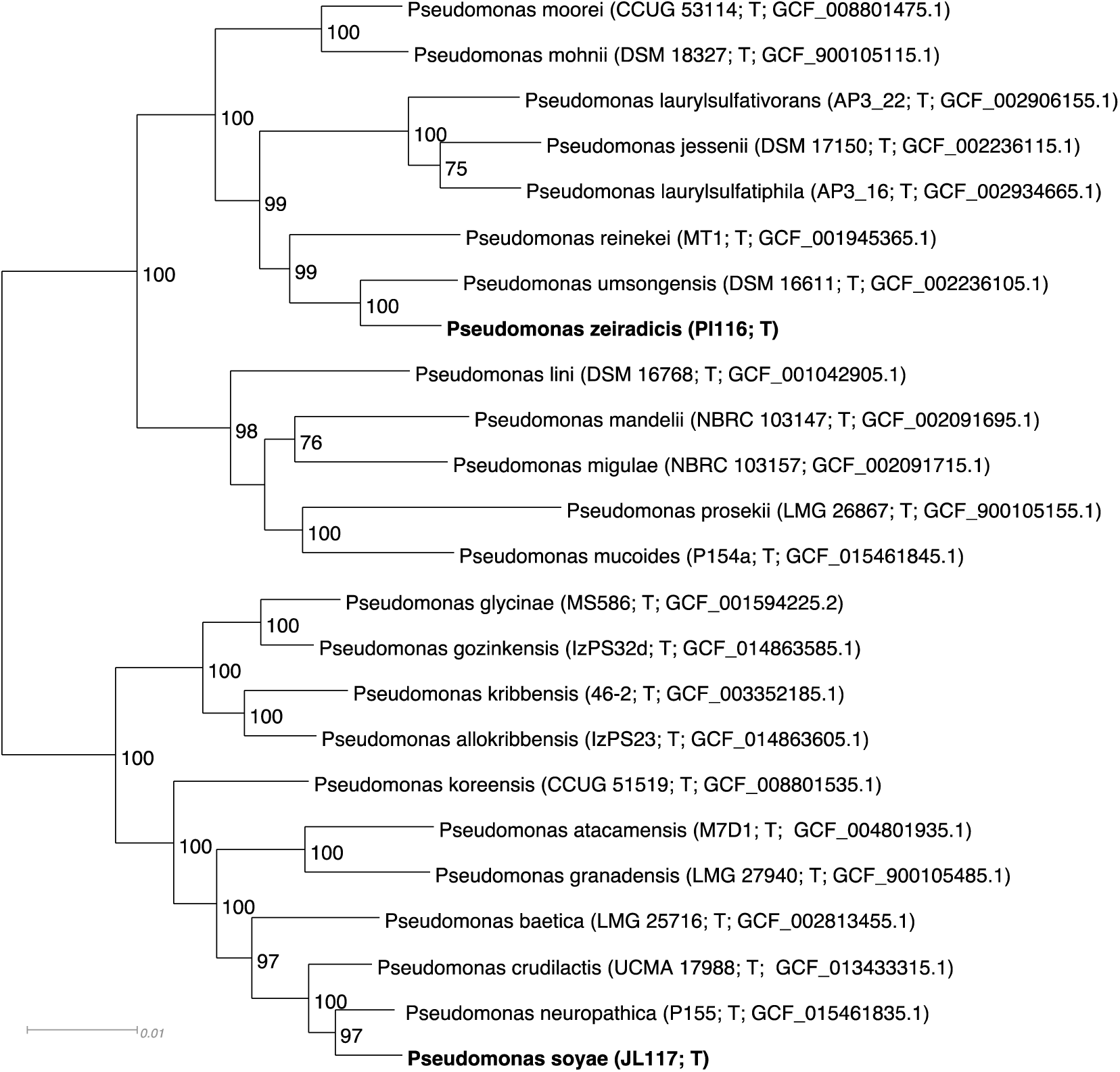
Phylogenomic tree of *Pseudomonas zeiradicis* sp. nov. strain PI116 and *Pseudomonas soyae* sp. nov. strain JL117 and relatives inferred by GToTree (Lee 2019). The species with the highest ANI to the new *Pseudomonas* species were included in the tree. Alignment consists of 38,231 amino acid positions from 160 – 170 genes depending on taxon. Species described in this study are in bold. Strain numbers and GenBank accession numbers follow species names, T stands for ‘type’. Support values above 50% are given by the branches. *Pseudomonas zeiradicis* sp. nov. strain PI116 is most closely related to *P. umsongensis* and *Pseudomonas soyae* sp. nov. strain JL117 to *P. neuropathica*, with 100% and 97% support, respectively. Branch lengths are proportional to the changes along the branches, a scale bar is provided at the bottom.

*Pseudomonas soyae* sp. nov. strain JL117

Strain JL117 shared 99.5% 16S rRNA gene sequence identity with *Pseudomonas reinekei* CCUG 53116^T^ and less with the remaining *Pseudomonas* species. A phylogenomic tree using GToTree (Lee 2019) confirmed the affiliation of strain JL117 with the genus *Pseudomonas*. JL117 was most closely related to *P. neuropathica* P155^T^ with 97% bootstrap support (Figure 1). The top average nucleotide identity (ANI) value of JL117 was 93.2% with *P. crudilactis* UCMA 17988^T^. This value was below the threshold for species demarcation (Richter and Rosselló-Móra 2009; Chun et al. 2018) providing further genomic support that strain JL117 represents a new genomic species of *Pseudomonas*.

### Geographic distribution and host range

Geographic distribution and host range of the novel species was inferred by comparison to congeneric genome assemblies of unidentified strains from GenBank. Hits to *Pseudomonas zeiradicis* sp. nov. strain PI116 included assembly GCF_016756945.1 from water of the western basin of Lake Erie (ANI: 99.2%; query coverage: 91.1%), assembly GCF_001006135.1 from urban soil in Iowa (ANI: 99.2%; query coverage: 91.3%), and assembly GCF_001976065.1 from compost in Massachusetts (ANI: 99.1%; query coverage: 92.6%). Hits to *Pseudomonas soyae* sp. nov. strain JL117 included assembly GCF_002003425.1 from the rhizosphere of *Glycine max* in Wisconsin (ANI: 96.2%; query coverage: 92.4%) and assembly GCF_003732405.1 from soil in Wyoming (ANI: 96.1%; query coverage: 93.3%). Known geographic distributions of the novel species from cultures is illustrated in Figure 2 and Figure 3, and substrates are compiled in Table 1.

**Figure 2.**
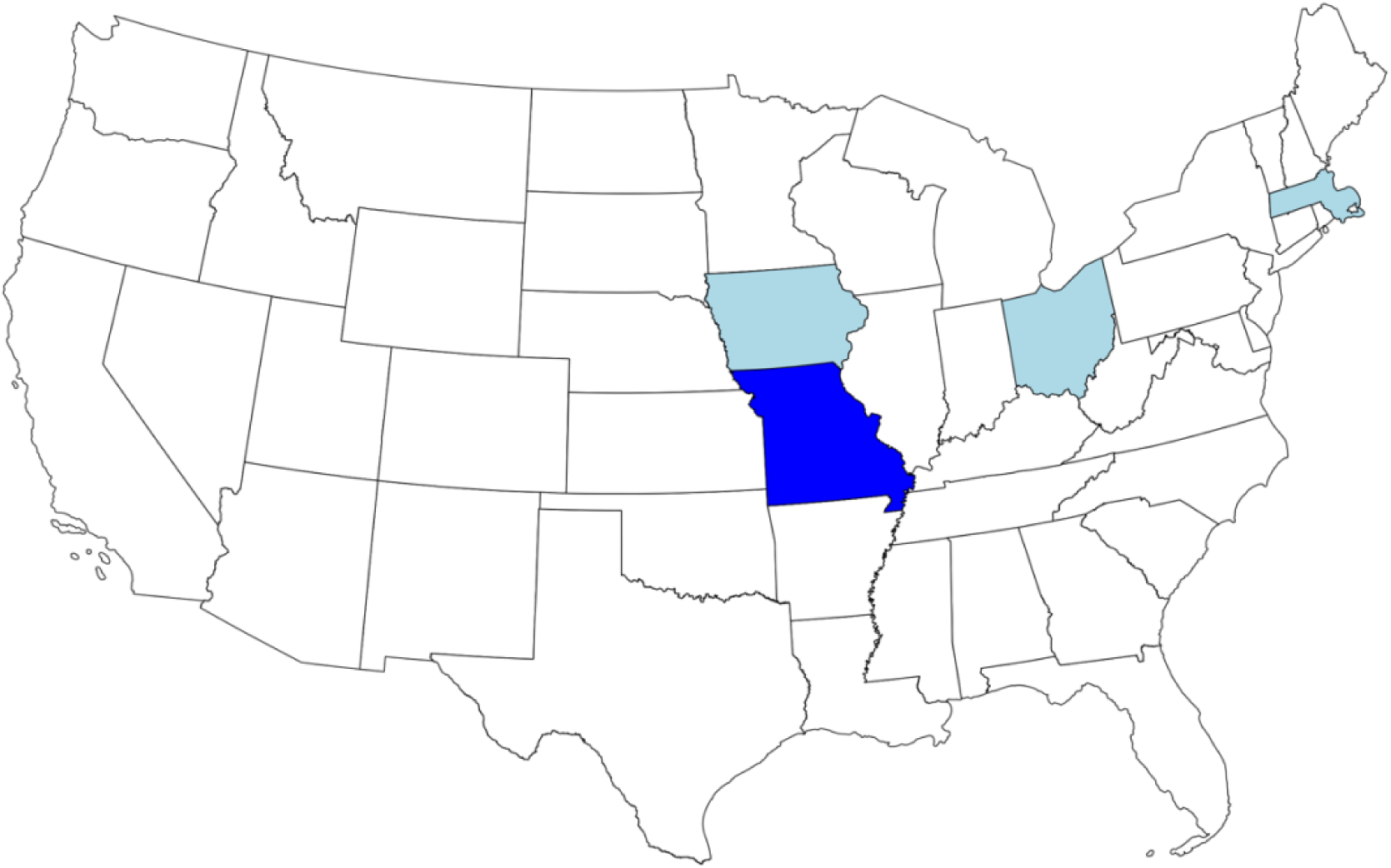
Geographic distribution of *Pseudomonas zeiradicis* sp. nov. strain PI116 based on culturing. Dark blue indicates state of origin for type strain, light blue are states of origin for additional conspecific strains based on GenBank genome assemblies. The strain isolated from Lake Erie water by the University of Toledo was assigned to Ohio. See text for details.

**Figure 3.**
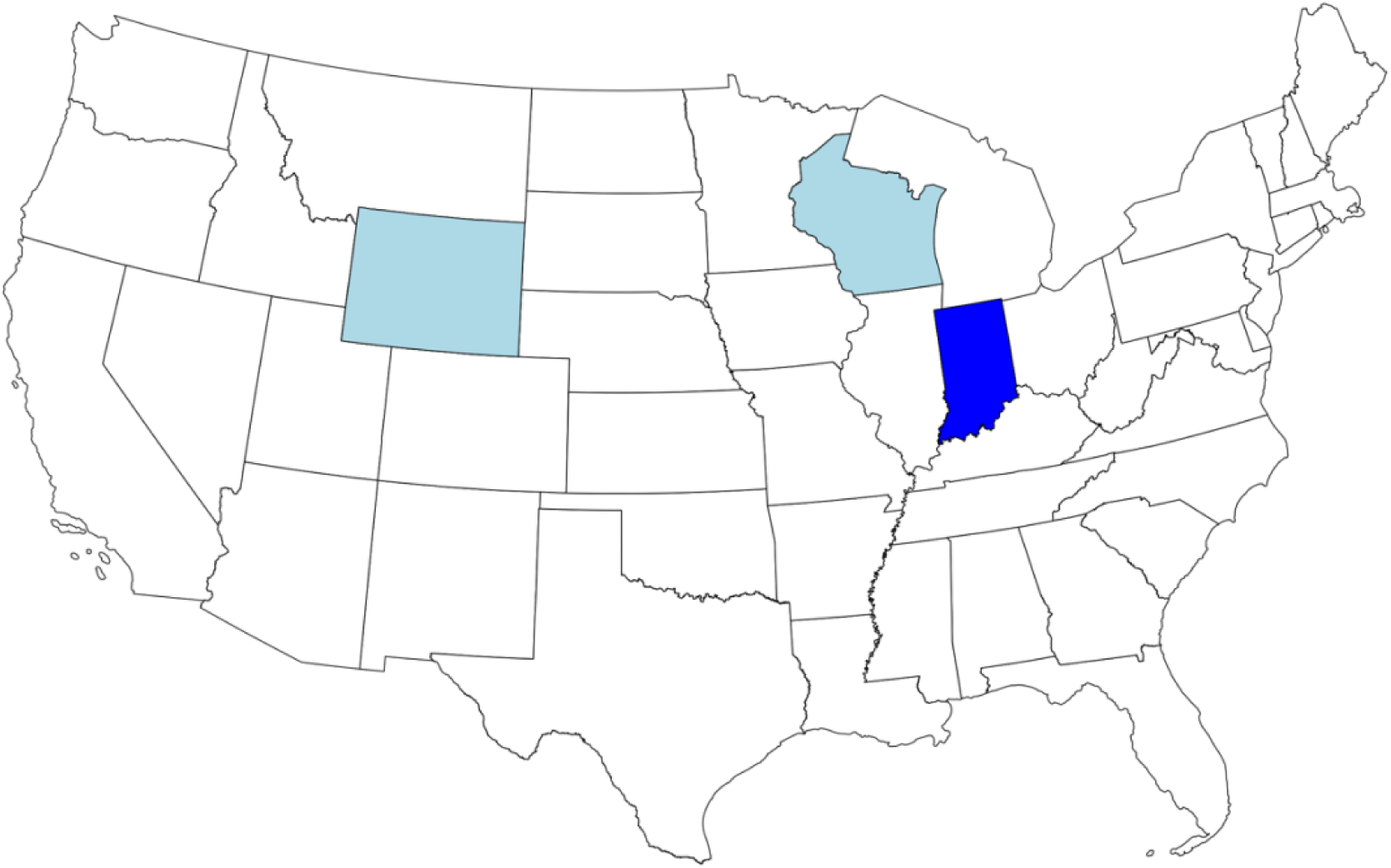
Geographic distribution of *Pseudomonas soyae* sp. nov. strain JL117 based on culturing. Dark blue indicates state of origin for type strain, light blue indicates states of origin for additional conspecific strains based on GenBank genome assemblies.

**Table 1.** Substrates of novel species based on culturing.

| Species | Substrate<sup>1</sup> |
| --- | --- |
| *Pseudomonas zeiradicis* sp. nov. strain PI116 | Compost, lake water, urban soil, *Zea mays* |
| *Pseudomonas soyae* sp. nov. strain JL117 | *Glycine max*, soil |

<sup>1</sup> No disease symptoms reported for strains isolated from plants.

**Table 2.** Physiological characteristics of newly described *Pseudomonas zeiradicis* sp. nov. strain PI116 and *P. soyae* sp. nov. strain JL117 and related *Pseudomonas* type strains.

|  | *P. zeiradicis*<br>PI116<sup>T</sup> | *P. soyae*<br>JL117<sup>T</sup> | *P. crudilactis*<br>UCMA<br>17988<sup>T</sup> | *P. neuropathica*<br>P155<sup>T</sup> | *P. umsongensis*<br>Ps 3-10<sup>T</sup> | *P. reinekei*<br>MT-1<sup>T</sup> |
| --- | --- | --- | --- | --- | --- | --- |
| General Characteristics: |  |  |  |  |  |  |
| Colony Morphology | Beige-yellow, circular, regular edges, mucoid | Beige-yellow, circular, regular edges, mucoid | Beige, circular, convex, regular edges, mucoid | Opaque, beige, circular, regular edges, mucoid | White-yellow, circular | White-yellow, circular, mucoid |
| Growth at 37°C | - | - | - | + | - | + |
| Growth at pH 5 | + | + | - | - | nr |  |
| Growth at pH 6 | + | + | + | - | nr |  |
| Enzymatic Activity: |  |  |  |  |  |  |
| Oxidase | + | + | + | + | + | + |
| Catalase | + | + | + | nr | + | nr |
| Urease | + | - | + | nr | - | - |
| Utilization of: |  |  |  |  |  |  |
| D-Maltose | - | - | + | - | - | - |
| D-Trehalose | - | + | - | + | - | - |
| Sucrose | - | + | - | - | - | - |
| N-Acetyl-D-Glucosamine | - | + | + | + | - | - |
| α-D-Glucose | + | + | + | + | + | + |
| Inosine | - | + | + | + | - | - |
| Glycerol | + | + | + | - | + | + |
| L-Arginine | + | + | + | + | nr | nr |
| L-Histidine | + | + | - | + | + | - |
| D-Galacturonic Acid | - | - | + | - | + | + |
| D-Glucuronic Acid | + | - | - | w | - | - |
| Glucuronamide | + | + | + | + | - | - |
| Formic Acid | + | + | + | - | + | + |
| Propionic Acid | + | + | - | + | + | w |
| Acetic Acid | + | + | - | + | + | + |
| α-Keto-Glutaric Acid | + | + | - | + | + | + |
| α-Keto-Butyric Acid | - | - | - | - | + | + |

+, Positive; w, Weakly positive; -, Negative; nr, Not reported.
Data from this study and Cámara et al. (2021), Duman et al. (2021), Kwon et al. (2003) and Schlusshuber et al. (2021).

### Morphology, physiology and biochemical characteristics

*Pseudomonas zeiradicis* sp. nov. strain PI116 and *Pseudomonas soyae* sp. nov. strain JL117

Strains PI116 and JL117 stained Gram-negative and had a rod shape (Figure 4, Figure 5). Growth was observed after 2 days on Tryptic soy (TSA) and R2A agars, with abundant biomass obtained after 5 days, at 30°C. No growth was observed at 37°C on any of the media tested. On TSA, colonies had a beige-yellow color, changing to opaque-white on R2A agar. Colonies were circular with regular edges, and smooth with a mucoid texture. An oxidative metabolism was recorded for both strains with growth at pH 5 and 6. In addition, the strains grew in 1 and 4% NaCl, but not at 8%. Catalase and oxidase were produced by the two bacteria, but urease activity was only observed for strain PI116. Physiological tests using Biolog GenIII Microplates showed that the new bacteria use a variety of organic compounds as carbon sources. These results are found in Table 1 and the species description.

**Figure 4.**
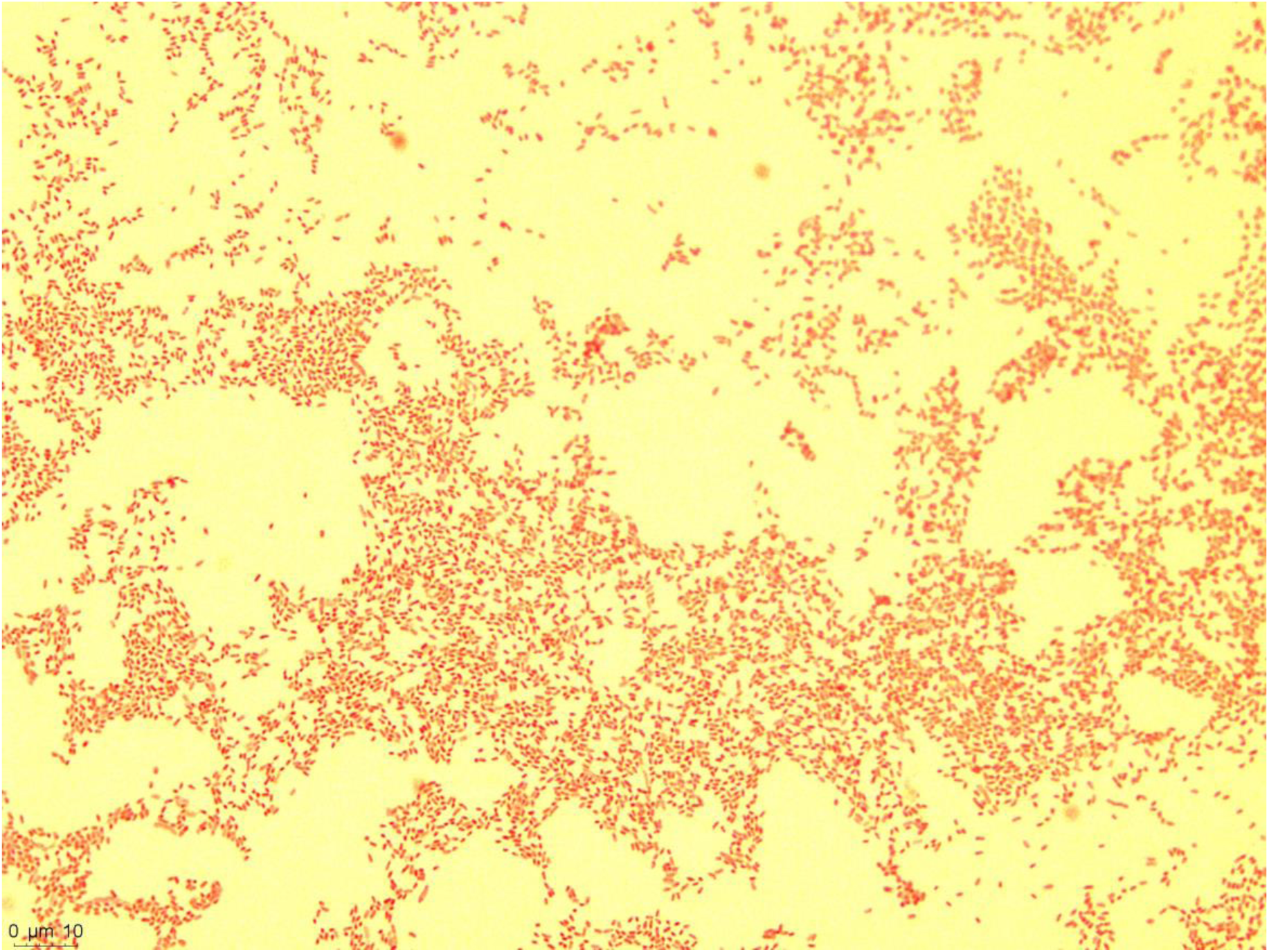
Morphology of *Pseudomonas zeiradicis* sp. nov. strain PI116 depicted following Gram stain using bright field microscopy. Bar = 10 µm.

**Figure 5.**
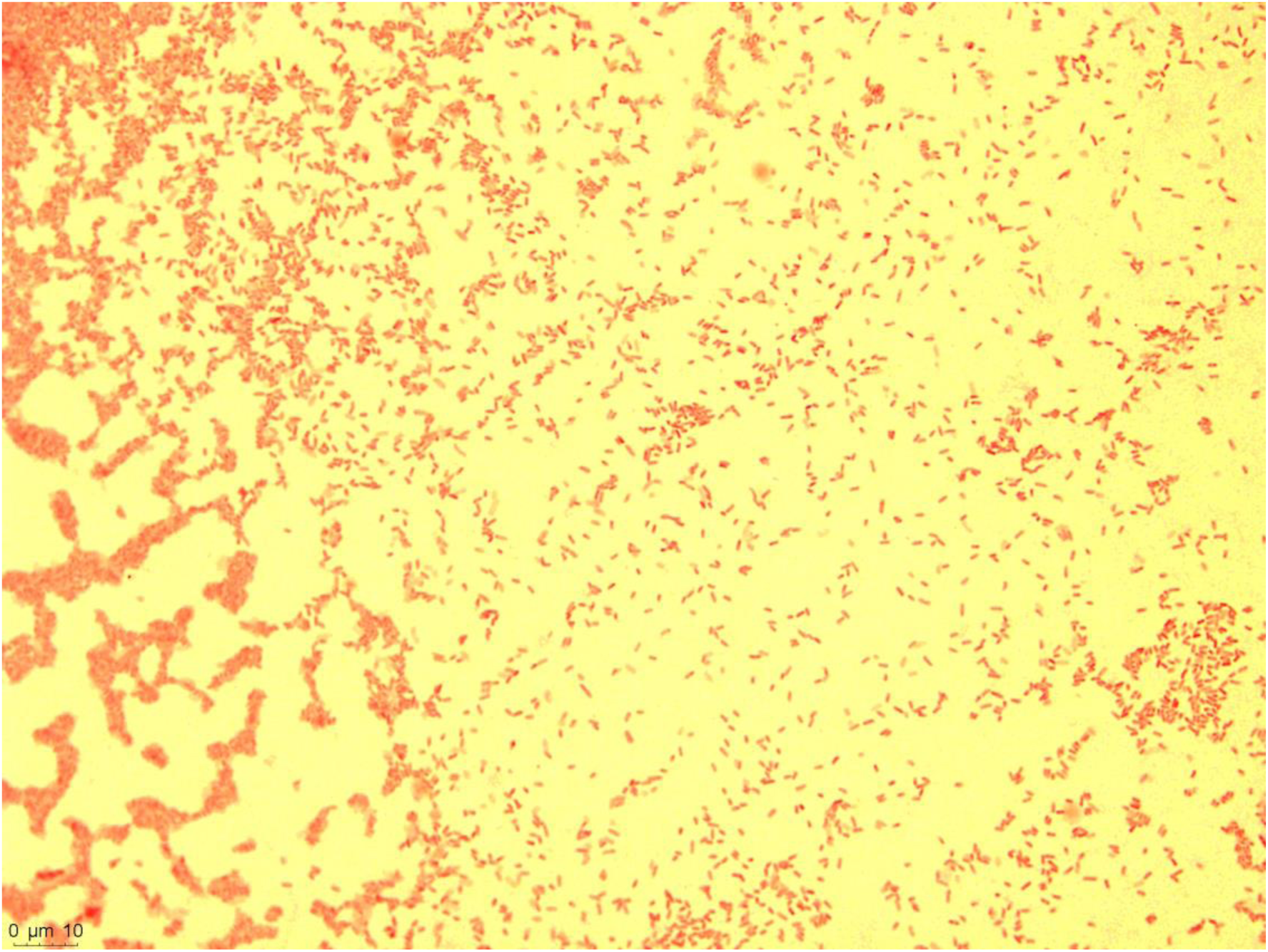
Morphology of *Pseudomonas soyae* sp. nov. strain JL117 depicted following Gram stain using bright field microscopy. Bar = 10 µm.

Strain PI116 showed resistance to lincomycin, troleandomycin, rifamycin RV, vancomycin, nalidixic acid and aztreonam, and susceptibility to minocycline according to Biolog GenIII microplate assays. The strain could grow in the presence of guanidine HCl, niaproof 4, but not in lithium chloride or potassium tellurite.

Production of urease and the utilization of D-galacturonic acid, D-glucuronic acid, glucuronamide and α-keto-butyric acid (Table 1) can be used to differentiate strain PI116 from its closest phylogenetic neighbor, *P. umsongensis* Ps 3-10^T^.

Similar to strain PI116, JL117 also had the same antibiotic resistance profile except that in addition to minocycline, it was susceptible to aztreonam. Growth of strain JL117 was inhibited by lithium chloride but not by potassium tellurite guanidine HCl or niaproof 4.

Strain JL117 can be differentiated from its closest phylogenetic neighbors *P. neurophatica* P155^T^ and *P. crudilactis* UCMA 17988^T^ by a combination of several physiological characteristics such as growth at 37°C, pH 5 and 6, and urease production. Several carbon source substrates that include trehalose, sucrose, glycerol, L-histidine and formic acid can also help to differentiate between strain JL117 and these species (Table 1).

### DESCRIPTION OF *PSEUDOMONAS ZEIRADICIS* SP. NOV. STRAIN PI116

*Pseudomonas zeiradicis* sp. nov. ze.i.ra’di.cis. L. fem. N. *zea* spelt, also the genus name of corn, *Zea mays;* L. fem. N. *radix* root; N.L. gen. n. *zeiradicis* of corn root Motile, short rod-shaped cells (1-2 μm long x 0.2-0.4 μm wide). Gram-stain negative. Colonies are opaque white to beige yellow, round, and mucoid. Aerobic and mesophilic. Catalase, oxidase, and urease positive. Growth is abundant at 30°C after 5 days on TSA and R2A agars, but not at 37°C. Growth is also observed at pH 5-7 and up to 4% NaCl. The following substrates are used as carbon sources: α-D-glucose, D-mannose, D-fructose, D-galactose, sodium lactate (1%), fusidic acid, DL-serine, D-mannitol, D-arabitol, glycerol, L-alanine, L-arginine, L-glutamic acid, L-pyroglutamic acid, D-glucuronic acid, D-gluconic acid, glucuronamide, mucic acid, quinic acid, D-saccharic acid, methyl pyruvate, L-lactic acid, citric acid, α-keto-glutaric acid, DL-malic acid, bromo-succinic acid, Tween 40, γ-amino-butyric acid, β-hydroxy-D,L-butyric acid, propionic acid, acetic acid and formic acid. Does not use the following sugars as carbon sources: dextrin, D-maltose, D-trehalose, D-cellobiose, gentibiose, sucrose, D-turanose, stachyose, D-raffinose, α-D-lactose, D-melibiose, β-methyl-D-glucoside. D-salicin, N-acetyl-D-glucosamine, N-acetyl-β-D-mannosamine, N-acetyl-D-galactosamine, N-acetyl neuraminic acid, 3-methyl-glucose, DL-fucose, L-rhamnose and inosine. Does not hydrolyze gelatin or pectin.

The type strain, PI116^T^, was isolated from the roots of healthy field-grown *Zea mays* in Missouri, United States.

### DESCRIPTION OF *PSEUDOMONAS SOYAE* SP. NOV. STRAIN JL117

*Pseudomonas soyae* sp. Nov., so’yae. N.L. gen. N. *soyae*, of soybean, referring to the isolation source of the strain

Motile, short rod-shaped cells (1-2 μm long x 0.2-0.4 μm wide). Colonies are opaque white to beige yellow, round, and mucoid. Gram-stain negative. Growth is abundant at 30°C after 5 days on TSA and R2A agars, but not at 37°C. Aerobic and mesophilic. Catalase and oxidase positive; urease negative. Grows at pH 5-7 and in the presence of 4% NaCl but not at 8%. The following substrates are used as carbon sources: D-trehalose, sucrose, N-acetyl-D-glucosamine, α-D-glucose, D-mannose, D-fructose, D-galactose, sodium lactate (1%), fusidic acid, DL-serine, D-mannitol, D-arabitol, glycerol, L-alanine, L-arginine, L-glutamic acid, L-pyroglutamic acid, D-gluconic acid, glucuronamide, mucic acid, quinic acid, D-saccharic acid, methyl pyruvate, L-lactic acid, citric acid, α-keto-glutaric acid, L-malic acid, Tween 40, γ-amino-butyric acid, β-hydroxy-D,L-butyric acid, propionic acid, and acetic acid. Does not use the following sugars as carbon sources: dextrin, D-maltose, D-cellobiose, gentibiose, D-turanose, stachyose, D-raffinose, α-D-lactose, D-melibiose, β-methyl-D-glucoside. D-salicin, N-acetyl-D-glucosamine, N-acetyl-β-D-mannosamine, N-acetyl-D-galactosamine, N-acetyl neuraminic acid, 3-methyl-glucose, DL-fucose, and L-rhamnose. Does not hydrolyze gelatin or pectin.

The type strain, JL117^T^, was isolated from *Glycine max* in Indiana, United States.

## ACKNOWLEDGEMENTS

We would like to thank Professor Aharon Oren, The Hebrew University of Jerusalem, for checking Latin species names.

